# A test of altitude-related variation in aerobic metabolism of Andean birds

**DOI:** 10.1101/2020.10.01.310128

**Authors:** Natalia Gutierrez-Pinto, Gustavo A. Londoño, Mark A. Chappell, Jay F. Storz

## Abstract

Endotherms at high altitude face the combined challenges of cold and hypoxia. Cold increases thermoregulatory costs, and hypoxia may limit both thermogenesis and aerobic exercise capacity. Consequently, in comparisons between closely related highland and lowland taxa, we might expect to observe consistent differences in basal metabolism (BMR), maximal metabolism (MMR), and aerobic scope. Broad-scale comparative studies of birds reveal no association between BMR and native elevation, and altitude effects on MMR have not been investigated. We tested for altitude-related variation in aerobic metabolism in 10 Andean passerines representing five pairs of closely related species with contrasting elevational ranges. Mass-corrected BMR and MMR were significantly higher in most highland species relative to their lowland counterparts, but there was no uniform elevational trend across all pairs of species.

**Summary statement:** We tested for altitude-related variation in aerobic metabolism in species pairs with contrasting elevational ranges. Metabolic rates were significantly higher in most highland species but there was no uniform elevational trend.

## Introduction

Endotherms that are native to high-altitude environments must contend with physiological challenges posed by the reduced partial pressure of O_2_ (*P*O_2_) and low ambient temperature (T_a_). Depending on acclimatization history, reduced *PO*_2_ may compromise the maximum capacities for aerobic exercise (MMR; maximum metabolic rate) due to the reduced availability of O_2_ to fuel ATP synthesis (Chappell et al., 2007; Hayes, 1989a; McClelland and Scott, 2019; Storz and Scott, 2019; Storz et al., 2010). BMR may be elevated in highland species due to increased thermoregulatory demands or due to a correlated response to changes in MMR (Hayes and Garland, 1995; Portugal et al., 2016; Rezende et al., 2004). Non-proportional changes in BMR and MMR entail changes in absolute aerobic scope, defined as the difference between the two rates (MMR – BMR), which reflects an animal’s capacity to increase its rate of aerobic metabolism above maintenance levels (Bennett, 1991; Hochachka, 1985).

In comparison with lowland relatives, mammals native to high-altitude often have higher mean MMR in hypoxia and suffer a smaller decrement in MMR with increasing hypoxia (Chappell and Dlugosz, 2009; Cheviron et al., 2012; Cheviron et al., 2014; Lau et al., 2017; Lui et al., 2015; Schippers et al., 2012; Storz et al., 2019; Tate et al., 2017; Tate et al., 2020). In addition, when measured at their native elevations, BMR is consistently higher in high-altitude deer mice *(Peromyscus maniculatus)* relative to lowland conspecifics (Hayes, 1989a; Hayes, 1989b), although it is not known to what extent the elevated BMR reflects an evolved change or a reversible acclimatization response. Available evidence for birds suggests that BMR does not vary with elevation, and it is unknown whether MMR exhibits a consistent pattern of altitudinal variation among species.

Studies of highland and lowland populations of rufous-collared sparrows *(Zonotrichia capensis)*, measured at their native altitudes, found no differences in BMR (Castro et al., 1985), thermogenic capacity (Novoa et al., 1990), or field metabolic rates (Novoa et al., 1991). A recent study involving a phylogenetically diverse set of more than 250 neotropical bird species found no significant association between BMR and native elevation (Londoño et al., 2015). However, altitude effects may be more readily detectable in fine-grained comparisons between pairs of closely related species that are native to different elevations but that are otherwise ecologically similar. Moreover, it remains unclear if exercise-induced MMR and aerobic scope exhibit consistent patterns of altitude variation.

We measured BMR, MMR, and aerobic scope in 10 Andean passerines, representing five pairs of closely related species with contrasting elevational ranges (Figure 1A, B). We used a paired lineage design (Felsenstein, 2004) such that the five pairwise comparisons were phylogenetically independent (Figure 1A).

**Figure 1.**
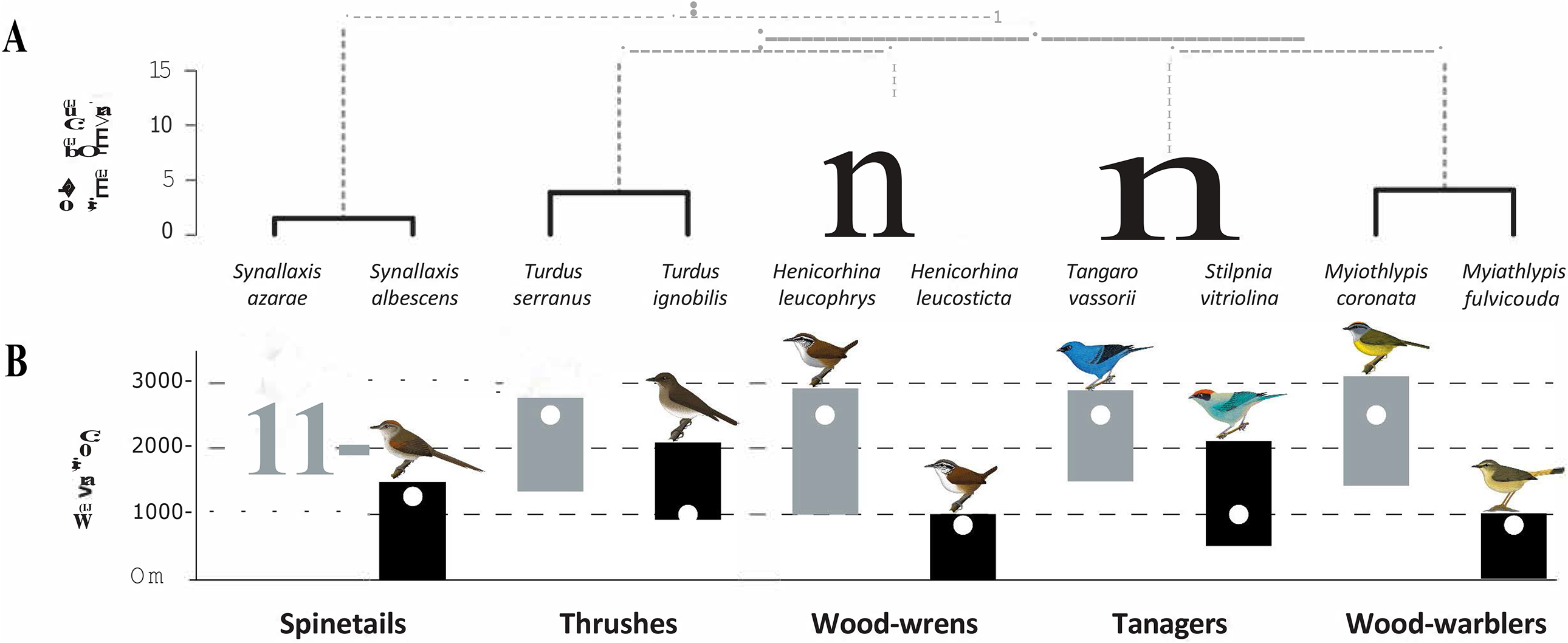
**A**. Phylogenetic relationships among the ten study species. Terminal branches (in black) connect pairs of high- and low-elevation species. Branch lengths are proportional to estimated divergence times (Barker et al., 2015; Batista et al., 2020; Cadena et al., 2019; Derryberry et al., 2011) **B.** Approximate elevational distribution of the study species in the northern Andes (Hilty and Brown, 1986). White circles represent the elevation at which measurements were made for each species (see supplementary table 1). Illustrations from Ayerbe-Quiñones (2018).

## Materials and Methods

### Experimental design

Birds were captured between June and August from 2017 to 2019 at several field sites in the western Andes of Colombia (figure 1A; supplementary Table 1). We compared closely related species that had contrasting elevation ranges (figure 1B) but which are otherwise similar ecologically (supplementary table 2). High elevation species were captured between 2300 and 2500 m; low elevation species were captured between 500 and 1400 m. Annual mean temperature differed between localities by approximately 6°C and ambient pO_2_ differed by 1.9 - 4.5 kPa (mean 3 kPa).

### Field protocol

We mist-netted birds (aided by playback of vocalizations) during the day (9:00 to 18:00 h). We released juveniles (identified on the basis of plumage or color of bill gape) and adults with brood patches. We measured MMR immediately after capture and subsequently kept birds inside cloth bags, taking them out every two hours to provide water and food, until 4 hours before the onset of BMR measurements (after sunset, around 19:00h), to ensure that the birds were post-absorptive (Karasov and del Rio, 2007). Body mass was measured immediately after capture and also before and after BMR measurements. All procedures were approved by the University of Nebraska IACUC (project ID 1499) and Colombian research permits granted to G.A.L. (Permit 536, May 20/2016).

### Respirometry procedures

We used a flow-through respirometry system to measure metabolic rates. Incurrent air was dried with silica gel and the flow was then divided into four metered channels using a FlowBar (Sable Systems, Las Vegas, NV, USA). One channel was used for reference air. The other three supplied air continuously to three metabolic chambers made of acrylic, each equipped with a thermocouple that measured excurrent air temperature. Another thermocouple measured T_a_ in the incubator. We measured one to three birds per night and matched incurrent airflow into chambers with the body mass of each tested bird (200–1000 ml min^-1^ STPD). Excurrent air flows were sampled sequentially by a multiplexer (Sable Systems RM-8). Subsamples of excurrent airflow (50–200 mL min^-1^) were scrubbed of CO_2_ and H_2_O (using soda lime and silica gel, respectively) and routed through a Sable Systems FoxBox to measure O_2_ content. Each bird was monitored for 15 min and reference air was measured for 2.5 min before switching between individuals; this pattern was repeated until the measurements were finished. T_a_ was kept constant (±0.5°C) using a PELT-5 controller (Sable Systems). Birds were first measured at T_a_ = 34°C and then at T_a_ = 30°C, remaining at each stable temperature for at least one hour. We recorded T_a_, flow rate and O_2_ content every second using Warthog LabHelper (www.warthog.ucr.edu) interfaced to a Sable U1-2 A-D converter.

We elicited MMR using forced exercise on a hop-flutter wheel (Chappell et al., 1999). This method reliably elicits behavioral exhaustion, with repeatable MMR (Chappell et al., 1996), but may not elicit maximum power output of the flight muscles. Accordingly, VO_2_ values measured with this technique are usually lower than those measured in wind tunnels (Chappell et al., 2011; McKechnie and Swanson, 2010). During exercise trials, incurrent air flow was 750 – 1000 mL min^-1^, and no multiplexer was used. The chamber was manually rotated until birds reached their maximum rates of oxygen consumption, typically at the beginning of the experiment. All tested birds attained similar levels of behavioral exhaustion during the exercise trials and no birds were injured during experiments.

### Data analysis

We calculated metabolic rates using LabAnalyst (www.warthog.ucr.edu). After baseline correction, we used the flow rate (FR; mL min^-1^ STPD), and the incurrent (FiO_2_; 0.2095) and excurrent (FeO_2_) oxygen concentrations to obtain *V*O_2_ (mL O_2_ min^-1^), applying the ‘Mode 1’ formula:

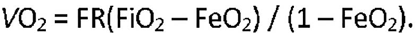

BMR was computed as the lowest continuous average *V*O_2_ over 5 min during periods of low and stable *V*O_2_, and the lowest value of the two temperature measurements per bird (30 and 34 °C) was chosen for subsequent analyses. Before obtaining MMR (*V*O_2max_), we applied the instantaneous correction (Bartholomew et al., 1981) to compensate for the mixing characteristics of the system (i.e., the blunted response to rapid changes in O_2_ concentration). We calculated MMRas the highest continuous averaged *V*O_2_ over 1 minute during periods of high and stable *V*O_2_ values. Finally, we calculated the absolute aerobic scope for each bird as the difference between MMRand BMR.

### Statistics

We used log-transformed metabolic rates in interspecific comparisons and we included log-transformed body mass as a covariate in the analyses. For each species and metabolic measurement, we discarded data points that fell outside ± 2 standard deviations from the mean, resulting in the removal of two data points for BMR, six for MMR, and five for aerobic scope.

To evaluate whether the allometric association between mass and *V*O_2_ differed between closely related species, we tried fitting standardized major axis (SMA) regressions between mass and *V*O_2_ for each species using the *smatr* package (Warton et al., 2012) for R v. 3.3.2 (R Core Team, 2012). Since most of the regressions were not significant (results not shown), we followed two different approaches to evaluate the influence of body mass on measured metabolic rates. First, we adjusted linear models to account for the joint influences of elevation and mass on VO_2_. For this, we fitted a linear mixed model (package *Ime4*; Bates et al., 2014) with VO_2_ as the response variable, elevation (categorical; 1 for high elevation, 2 for low elevation) as a fixed effect, and mass (continuous) per species pair (categorical; 5 levels) as a random effect:

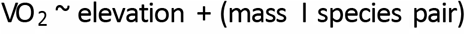

This model allows the slope of the relationship between mass and VO_2_ to be different for each species group. To further explore whether the effect of elevation differs between species pairs after accounting for mass, we also ran a nested ANCOVA, where the variation in VO_2_ is explained by mass and by elevation per group:

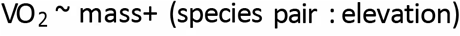

Second, in order to better understand the trends observed in the linear models, we mass-corrected each of our observed *V*O_2_ values (BMR, MMR, aerobic scope) by dividing them by M_b_^S^ (Gillooly et al., 2001), where M_b_ is body mass for each individual and S is the allometric scaling coefficient obtained by McKechnie and Wolf (2004) for passerine birds (S = 0.667). We then used t-tests to evaluate the statistical significance of the difference in mass-corrected *V*O_2_ between high- and low-elevation species within each pair. We also ran a *post-hoc* power analysis (package *pwr* v1.3; Champely, 2020) using Hedges’ G (Hedges, 1983) to estimate effect sizes with a significance level of 0.05.

Finally, to assess the influence of other variables on metabolic rates, we built linear mixed models that included various combinations of mass and the interaction between species group (categorical; 5 levels) and elevation (categorical; high, low) as fixed factors, and age (categorical; immature, adult), sex (categorical; male, female), molt state (categorical; absent if bird had no developing feathers, moderate if few, abundant if several), and year of capture (2017, 2018, 2019) as random factors. In all mixed effect models the variance explained by any variable other than mass was negligible (usually ≪ 0.1), and the model that included mass as the only predictor had the lowest AIC (ΔAIC = 46; results not shown).

## Results and Discussion

We measured 96 wild-caught birds, with sample sizes of 8–12 individuals per species (Figure 2; supplementary table 3), except for *Turdus serranus* (n=4). Both the linear mixed model and the ANCOVA explained a high percentage of variation in the data (average R^2^ for mixed model: 0.98; average adjusted R^2^ for ANCOVA: 0.69; supplementary tables 4 and 5). Elevation was correlated with BMR, MMR, and aerobic scope, but only after accounting for the effects of species pair and body mass, as evidenced by the difference between the average marginal R^2^ (0.006; variance explained by the fixed effect) and the average conditional R^2^ (0.98; variance explained by the full model) (supplementary table 4). Likewise, after accounting for mass, elevation also had a significant effect on variation in BMR, MMR, and aerobic scope in each species pair (supplementary table 5), with the exception of BMR in thrushes. In all species pairs other than thrushes, high-elevation species had higher metabolic rates than their lowland counterparts, as indicated by the negative slopes estimated for the effect of elevation on each species group (supplementary table 5).

**Figure 2.**
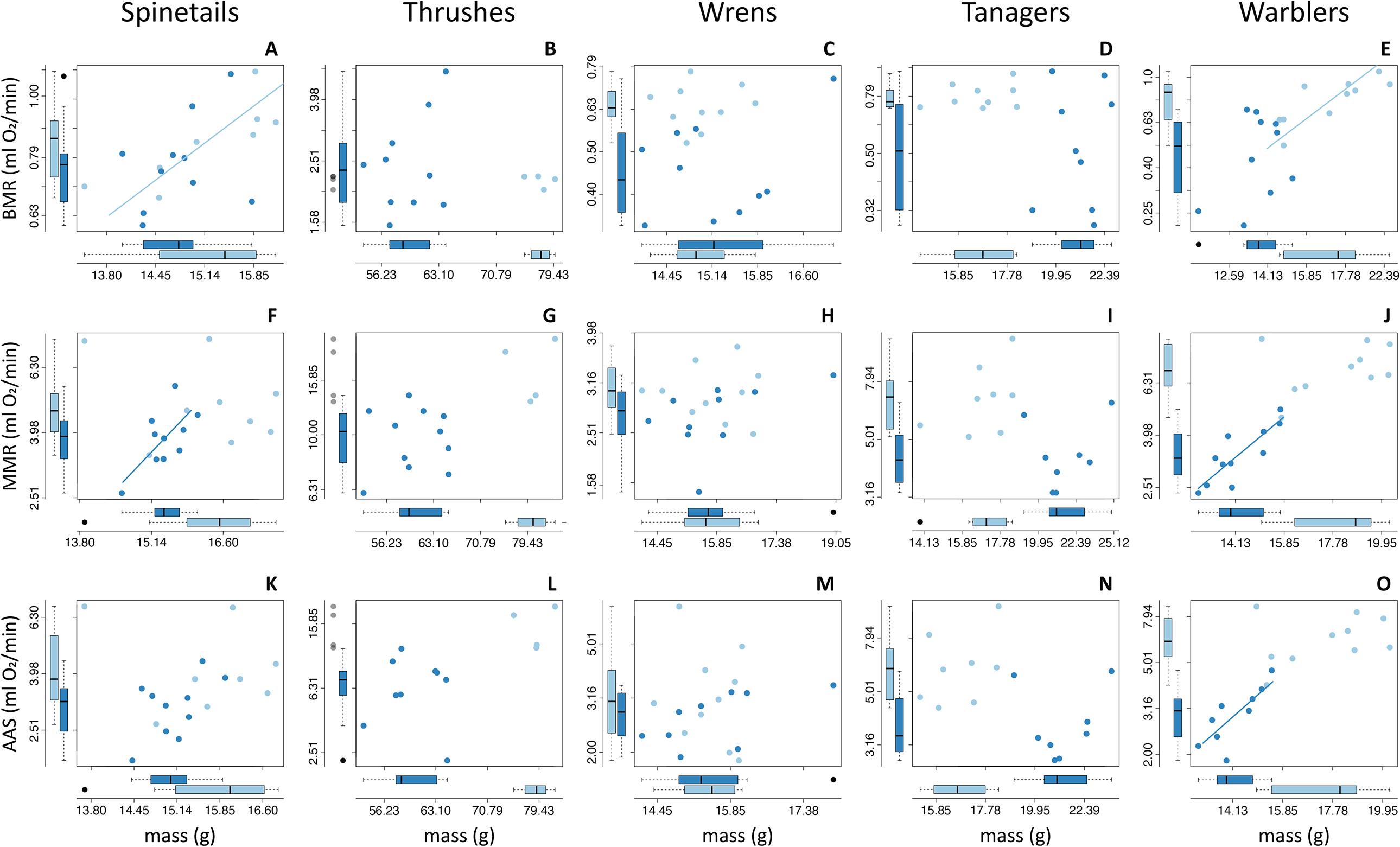
Relationships between mass and BMR (top row), MMR (middle row), and absolute aerobic scope (AAS; bottom row) for each pair of high- and low-elevation species. SMA regression lines are shown in cases where the tested association was statistically significant. In each pair, high elevation species are shown in light blue and low elevation species in dark blue. Boxplots depict the variation for each species in metabolic rate (left) and mass (bottom).

Our pairwise comparisons of mass-corrected metabolic rates revealed significant differences between high- and low-elevation species in most but not all cases. BMR was significantly higher in high-elevation wrens, tanagers, and warblers (Figure 3, left; supplementary table 3), MMR was significantly higher for high-elevation wrens, tanagers and warblers (Figure 3, middle), and aerobic scope was significantly higher for high-elevation thrushes, tanagers, and warblers (Figure 3, right). Spinetails did not exhibit significant elevational differences in BMR, MMR, or aerobic scope.

**Figure 3.**
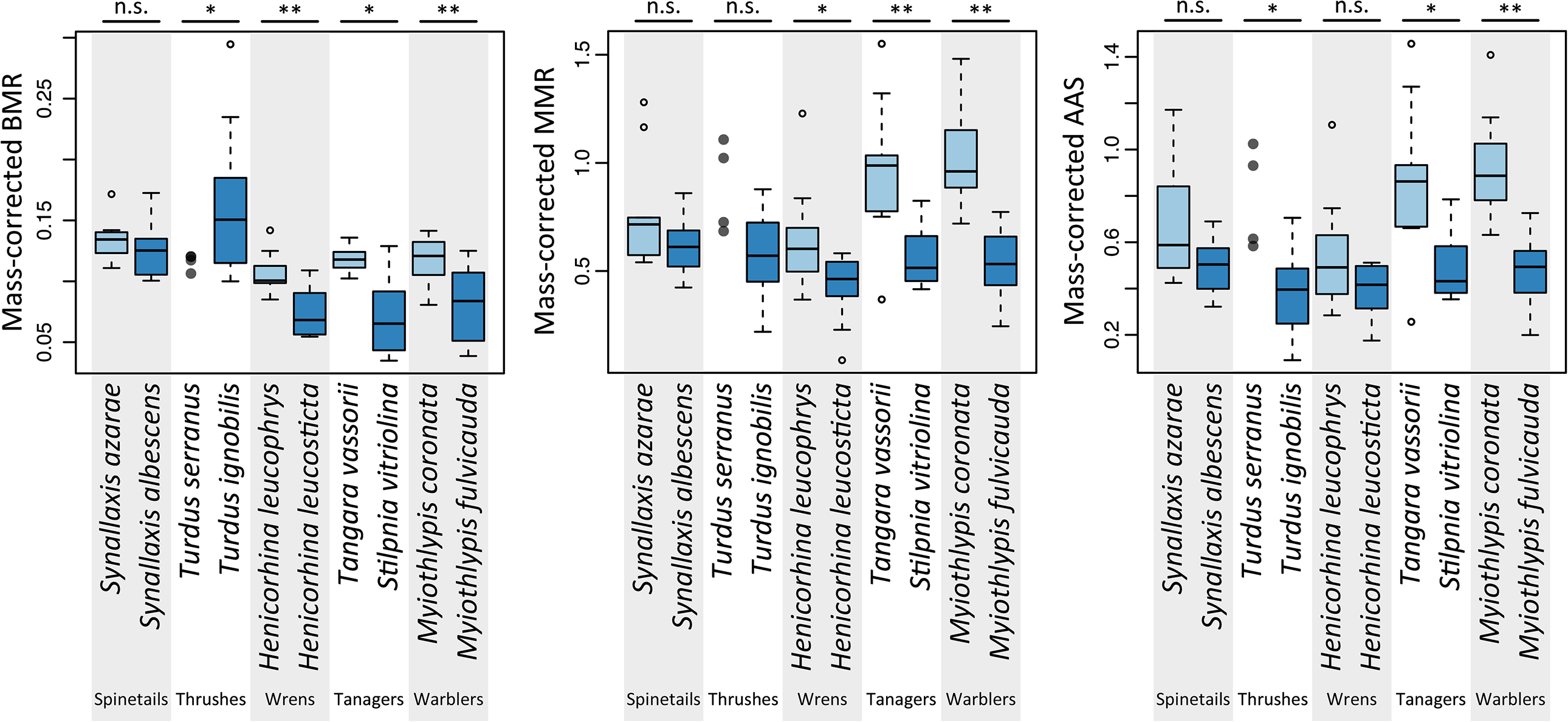
Mass-corrected values (ml O_2_ min^-1^ g^-1^) for BMR (top), MMR (middle), and absolute aerobic scope (AAS; bottom) of Andean passerines. Alternating light grey and white columns denote the species pairs being compared; within each pair, light and dark blue boxes represent high and low elevation species, respectively. Three significance levels of the pairwise t-tests comparing mass-corrected metabolic values between high- and low-elevation species are indicated on top of each graph: non-significance (n.s; *p* > 0.05), *p* < 0.05 (*), and *p* < 0.01 (**); *p-*values and differences in group means can be found in supplementary table 3.

In our small sample of Andean passerines, we found a significant effect of elevation on BMR, MMR, and aerobic scope after accounting for effects of body mass (supplementary tables 3, 4, and 5). In most cases, high-elevation species had higher BMR and MMR than closely related and ecologically similar low-elevation species. Overall, we observed significant differences in rates of aerobic metabolism between individual pairs of species, but we did not document a uniformly consistent elevational trend. Our study and most others to date have investigated elevational variation in aerobic metabolism by measuring wild-caught birds in their native habitat (e.g. Jones et al., 2020; Londoño et al., 2015; Londoño et al., 2017). In the future, common-garden or reciprocal-transplant experiments that control for acclimatization effects should help reveal whether bird species native to high elevations have generally evolved increased aerobic performance capacities in cold, hypoxic conditions.

## Acknowledgements

We wish to thank all the students whose invaluable help in the field made this project possible, especially to Jorge Lizarazo, Isabel Cifuentes, Valentina Echeverry, Jose Riascos, and Maria Laura Mahecha. For facilitating logistics at fieldwork sites in Colombia we thank the staff at ICESI University, staff at CELSIA, Alirio Bolivar, Ana Tulia Montes, and Gustavo Giraldo. Additional help with fieldwork preparation was provided by Andrew Crawford and Santiago Herrera. Gwen Bachman offered vital guidance while processing metabolic rate data. We are thankful with Kate Lyons, John DeLong, Graham Scott, and Grant McClelland for discussions and assistance with the statistical analyses. Finally, this work greatly benefited from discussions with Gwen Bachman, Chris Witt, Kristi Montooth, Colin Meiklejohn, and Jamilynn Poletto.

## Author contributions

Conceptualization: N.G.P., J.F.S.; Methodology: N.G.P., J.F.S.; Formal analysis: N.G.P.; Investigation: N.G.P.; Data curation: N.G.P.; Writing – original draft: N.G.P.; Writing – review & editing: N.G.P, G.A.L., M.A.C., J.F.S.; Supervision: G.A.L., J.F.S.; Funding acquisition: N.G.P, G.A.L., J.F.S.

## Supplementary Tables

**Supplementary table 1.** Information about geographical coordinates, elevation (m), weather, and landscape matrix of the localities visited for this study. All localities visited are located in the Valle del Cauca department, Colombia. Annual mean temperature and precipitation were extracted from the WorldClim database (BIO1 and BIO12, respectively; Hijmans et al, 2005^*a*^). Also included are the species captured in each site and the year(s) each locality was visited.

| Name | Coordinates | Elevation (m) | Annual Mean Temperature (°C) | Annual precipitation (mm) | Ambient pO2 (kPa) | Landscape matrix | Species captured | Years visited |
| --- | --- | --- | --- | --- | --- | --- | --- | --- |
| Estación Biológica Zygia-ICESI | 3.442°N, 76.662°W | 2311-2553 | 17.5 | 2260 | 15.6-16.1 | Primary and secondary forests | *Synallaxis azarae*<br>*Turdus serranus*<br>*Henicorhina leucophrys*<br>*Tangara vassorii*<br>*Myiothlypis coronata* | 2017-2019 |
| Alto Anchicayá | 3.576°N, 76.880°W | 525-889 | 24.2 | 2093 | 19.2-20.0 | Primary forests and road sides | *Henicorhina leucosticta*<br>*Myiothlypis fulvicauda* | 2017-2018 |
| Universidad ICESI | 3.341°N, 76.529°W | 1014-1021 | 23.6 | 1490 | 18.9 | Urban university campus | *Turdus ignobilis*<br>*Tangara vitriolina* | 2017-2018 |
| Ecoparque el Embudo | 3.345°N, 76.553°W | 1099 | 23.2 | 1557 | 18.7 | Secondary forests and cleared areas | *Tangara vitriolina* | 2019 |
| Finca el Pasatiempo | 4.677°N, 75.739°W | 1420-1445 | 20.4 | 2251 | 17.9 | Coffee farms and road sides | *Synallaxis albescens* | 2019 |
| Finca el Tabacal | 4.681°N, 75.764°W | 1266-1276 | 21.3 | 1973 | 18.3 | Coffee farms and road sides | *Synallaxis albescens* | 2019 |
| Rural Alcala | 4.676°N, 75.772°W | 1276 | 21.4 | 1927 | 18.3 | Coffee farms and road sides | *Synallaxis albescens* | 2019 |
| Finca Tipitapa | 4.662°N,<br>75.761°W | 1334 | 21.3 | 1965 | 18.2 | Coffee farms and<br>road sides | *Synallaxis albescens* | 2019 |
| Finca Villa Ramona | 4.661°N,<br>75.754°W | 1337 | 21.0 | 2052 | 18.2 | Coffee farms and<br>road sides | *Synallaxis albescens* | 2019 |

<sup>a</sup>Hijmans, R. J., Cameron, S. E., Parra, J. L., Jones, P. G. and Jarvis, A. (2005). Very high resolution interpolated climate surfaces for global land areas. *International Journal of Climatology* **25**, 1965-1978.

**Supplementary table 2.** General information of the biology of the species selected for this study. We paired closely related species that had contrasting elevation ranges but were similar in terms of morphology, life history, and foraging ecology.

| Species | Size | Habitat | Diet | Foraging | Breeding | Behavior | Sources |
| --- | --- | --- | --- | --- | --- | --- | --- |
| Azara's Spinetail<br>*Synallaxis azarae* | 15-18 cm,<br>12-18 g. | Dense undergrowth of forest edges secondary forest, and overgrown roadsides, usually above 1600 m. | Mostly arthropods, occasionally small seeds. | Captures prey from foliage, small branches and dead leaves, within 1–2 m of ground. | Throughout the year in Colombia. | Sedentary and territorial, usually in pairs, rarely joining mixed species flocks. | Hilty and Brown (1986); Remsen (2020a) |
| Pale-breasted Spinetail<br>*Synallaxis albescens* | 13-16 cm,<br>9-17 g. | Open areas with grassland, second growth scrub and overgrown roadsides, usually below 1500 m. | Mostly invertebrates. | Captures prey from foliage, grass and small branches, within 1–2 m of ground. | February to November in Colombia. | Sedentary and territorial, usually in pairs. | Hilty and Brown (1986); Remsen (2020b) |
| Glossy-black Thrush<br>*Turdus serranus* | 23-25 cm,<br>70-90 g. | Primary humid montane forest, forest borders, occasionally in open areas, mainly within 1400 to 2800 m. | Fruits and berries. | Mainly forages arboreally, sometimes along roads. | March to August in Colombia. | Sedentary, usually single or in pairs, may gather at fruiting trees. Rarely in mixed-species flocks. | Hilty and Brown (1986); Collar (2020) |
| Black-billed Thrush<br>*Turdus ignobilis* | 18-24 cm,<br>48-81 g. | Semi-open to open areas, forest borders, mainly within 900 to 2100 m. | Mainly invertebrates and fruits, occasionally seeds. | Mainly forages arboreally, also often on ground, usually near or in forests. | December to August in Colombia. | Sedentary, usually single or in pairs, may gather at fruiting trees. | Hilty and Brown (1986); Collar et al (2020) |
| Gray-breasted Wood-wren<br>*Henicorhina leucophrys* | 10-11 cm,<br>16-17 g. | Undergrowth of humid mountain forest, usually above 1500 m. | Mostly invertebrates. | Captures prey in dense vegetation, usually below 2 m. | December to June in Colombia. | Sedentary and territorial, usually in pairs. | Hilty and Brown (1986); Kroodsma et al (2020) |
| White-breasted Wood-wren<br>*Henicorhina leucosticta* | 10-11 cm,<br>15 g. | Undergrowth of wet lowland forest, from sea-level up to 1300 m. | Mostly invertebrates. | Captures prey in tangles around fallen trees and dense vegetation, from the ground up to 2–3 m. | January to July in most of its range. | Sedentary and territorial, usually in pairs, rarely joining ant follower flocks. | Hilty and Brown (1986); Kroodsma and Brewer (2020) |
| Blue-and-black Tanager<br>*Tangara vassorii* | 14-18 cm,<br>15-21 g. | Primary and secondary humid montane forest, and forest borders, from 1500 to 3500 m. | Mostly fruits and some insects. | Feeds in all forest levels but prefers the canopy, especially around fruiting trees. | February to August in Colombia. | Very active, usually seen in pairs or small groups, and often joins large mixed-species flocks. | Hilty and Brown (1986); Bernabe and Burns (2020) |
| Scrub Tanager<br>*Stilpnia vitriolina* | 14 cm,<br>18-26 g | Scrubby or open areas with sparse trees and bushes, mostly between 500 to 2200 m. | Mostly fruits and arthropods. | Feeds in all forest levels but prefers the canopy, especially around fruiting trees. | Throughout the year in Colombia. | Very active, usually seen alone, in pairs or small groups, rarely found with mixed-species flocks. | Hilty and Brown (1986); Hilty (2020) |
| Russet-crowned Warbler<br>*Myiothlypis coronata* | 14 cm,<br>13-19 g. | Humid montane forests, forest borders, and secondary growth, between 1300 to 2500 m. | Mostly invertebrates. | Captures prey at low to middle levels and dense areas, mainly at 1 to 6 m but can be seen higher. | May to June in Colombia. | Very active, usually seen in pairs or small groups, and often joins mixed-species flocks. | Hilty and Brown (1986); Curson and Bonan (2020); personal observations |
| Buff-rumped Warbler<br>*Myiothlypis fulvicauda* | 13 cm,<br>14 g | Primary and secondary forest areas, mostly around water bodies and forest edges. Usually below 1000 m. | Mostly invertebrates. | Captures prey close to the ground, along stream edges and damp areas, but can be seen higher. | February to April in Colombia. | Very active, usually seen in pairs. | Hilty and Brown (1986); Curson (2020); personal observations |

**Supplementary table 3.** Average mass-corrected BMR, MMR, and aerobic scope values, standard deviations (SD) and sample size (n) for each species analyzed. Also shown, the percentage of change in the highland mass-corrected metabolic rates in relation to lowland (positive values indicate higher values in the highland species), the respective pairwise t-tests evaluating the difference in means, the size of the effect, and the power of each statistical comparison.

|  | High elevation species |  |  | Low elevation species |  |  | % change | t-test | Power analysis |  |
| --- | --- | --- | --- | --- | --- | --- | --- | --- | --- | --- |
|  | Average<br>(mlO<sub>2</sub> min<sup>-1</sup>g<sup>-1</sup>) | SD | *n* | Average<br>(mlO<sub>2</sub> min<sup>-1</sup>g<sup>-1</sup>) | SD | *n* |  | *p*-value | effect size<br>(Hedges' G) | power |
| BMR |  |  |  |  |  |  |  |  |  |  |
| Spinetails | 0.135 | 0.018 | 8 | 0.127 | 0.023 | 10 | 6.3% | 0.4239 | 0.378 | 0.197 |
| Thrushes | 0.114 | 0.006 | 4 | 0.162 | 0.062 | 10 | -29.5% | **0.0383** | 0.888 | 0.619 |
| Wrens | 0.107 | 0.016 | 11 | 0.074 | 0.019 | 10 | 44.4% | **0.0005** | 1.857 | 1.000 |
| Tanagers | 0.119 | 0.011 | 9 | 0.074 | 0.034 | 9 | 60.4% | **0.0039** | 1.770 | 0.999 |
| Warblers | 0.117 | 0.019 | 10 | 0.082 | 0.032 | 10 | 43.5% | **0.0085** | 1.357 | 0.987 |
| MMR |  |  |  |  |  |  |  |  |  |  |
| Spinetails | 0.771 | 0.269 | 9 | 0.610 | 0.120 | 10 | 26.5% | 0.1265 | 0.789 | 0.658 |
| Thrushes | 0.896 | 0.230 | 4 | 0.587 | 0.191 | 12 | 52.5% | 0.0661 | 1.540 | 0.988 |
| Wrens | 0.637 | 0.241 | 11 | 0.430 | 0.155 | 11 | 48.1% | **0.0283** | 1.022 | 0.912 |
| Tanagers | 0.974 | 0.339 | 9 | 0.563 | 0.152 | 8 | 73.1% | **0.0069** | 1.534 | 0.991 |
| Warblers | 1.024 | 0.219 | 10 | 0.538 | 0.144 | 12 | 90.3% | **0.0000** | 2.674 | 1.000 |
| AAS |  |  |  |  |  |  |  |  |  |  |
| Spinetails | 0.679 | 0.276 | 8 | 0.493 | 0.112 | 10 | 37.7% | 0.1071 | 0.926 | 0.770 |
| Thrushes | 0.794 | 0.230 | 4 | 0.389 | 0.193 | 10 | 103.8% | **0.0283** | 1.989 | 0.999 |
| Wrens | 0.542 | 0.244 | 10 | 0.398 | 0.109 | 10 | 36.3% | 0.1122 | 0.764 | 0.654 |
| Tanagers | 0.865 | 0.351 | 9 | 0.491 | 0.161 | 8 | 76.1% | **0.0146** | 1.339 | 0.966 |
| Warblers | 0.927 | 0.224 | 10 | 0.474 | 0.153 | 10 | 95.5% | **0.0001** | 2.363 | 1.000 |

**Supplementary table 4.** Results of the linear mixed models that explained the variation in each metabolic parameter (logBMR, log MMR, and log AAS) as a function of elevation (fixed effect) and the interaction between species group and mass (random effects).

| *Predictors* | **BMR** |  |  | **MMR** |  |  | **AAS** |  |  |
| --- | --- | --- | --- | --- | --- | --- | --- | --- | --- |
|  | *Estimates* | *CI* | *p* | *Estimates* | *CI* | *p* | *Estimates* | *CI* | *p* |
| (Intercept) | -0.11 | -0.19 – -0.04 | **0.004** | 0.73 | 0.63 – 0.82 | **<0.001** | 0.65 | 0.56 – 0.73 | **<0.001** |
| Elevation | -0.07 | -0.13 – -0.01 | **0.022** | -0.15 | -0.20 – -0.10 | **<0.001** | -0.2 | -0.26 – -0.13 | **<0.001** |
| **Random Effects** | *Variance* |  |  | *Variance* |  |  | *Variance* |  |  |
| Residuals | 0.02 |  |  | 0.01 |  |  | 0.02 |  |  |
| Species group | 1.76 |  |  | 1.41 |  |  | 0.66 |  |  |
| Mass | 1.16 |  |  | 0.95 |  |  | 0.49 |  |  |
| Sample size | 90 |  |  | 90 |  |  | 84 |  |  |
| Marginal R<sup>2</sup>/<br>Conditional R<sup>2</sup> | 0.001 / 0.991 |  |  | 0.004 / 0.992 |  |  | 0.013 / 0.970 |  |  |

**Supplementary table 5.**
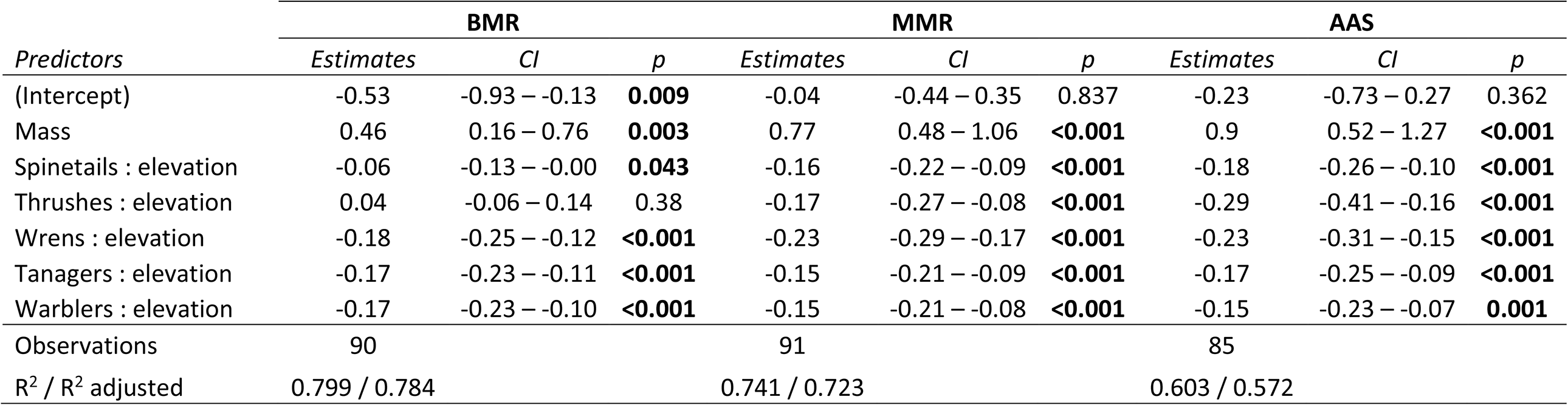
Results of the ANCOVA that explained the variation in each metabolic parameters (logBMR, log MMR, and log AAS) as a function of mass, and elevation per group.

